# Associations between absolute neutrophil count and lymphocyte-predominant breast cancer

**DOI:** 10.1101/551465

**Authors:** Chang Ik Yoon, So Eun Park, Yoon Jin Cha, Soong June Bae, Chi Hwan Cha, Da Young Lee, Sung Gwe Ahn, Joon Jeong

## Abstract

Tumor-infiltrating lymphocytes (TILs) might be associated with host-cell mediated immunity, which could be partly reflected by peripheral blood cell counts. We aimed to investigate whether peripheral blood cell counts are associated with TILs in breast cancer. Between August 2016 and July 2018, we evaluated the percentage of stromal TILs in breast cancer patients who underwent primary surgery, using the standardized methodology proposed by the international TIL Working Group. Lymphocyte-predominant breast cancer (LPBC) was defined as tumors having high TIL levels (≥ 50%). Peripheral blood cell counts including absolute neutrophil counts (ANC), absolute lymphocyte counts (ALC) and neutrophil-to-lymphocyte ratio (NLR) was obtained from pretreatment laboratory data. Of the 684 patients, 99 (17.2%) had LPBC, and 478 (82.8%) had non-LBPC. In a comparison of 3 markers of peripheral blood counts, LPBC had a significantly lower mean ANC than non-LPBC (3,330 vs. 3,660; *P*=0.004), but the other means were not different. Decreasing ANC was an independent clinical factor in predicting LPBC (OR: 0.736, 95% CI: 0.591-0.917; *P*=0.004). Low peripheral ANC might be linked with LPBC, supporting the hypothesis that systemic immune cell counts might be associated with the tumor-immune microenvironment.

## I. Introduction

Leukocytes, or white blood cells (WBCs), are involved in both immune and inflammatory processes. There are two primary types of leukocytes: granular and agranular. Granular leukocytes include eosinophils, neutrophils, and basophils, while agranular leukocytes, which lack cytoplasmic granules, include monocytes and lymphocytes. In general, leukocytes play a major role in the immune response to macromolecules released from abnormal cells and microorganisms.

Increasing evidence has shown that blood cell counts or ratio, such as the WBC count, absolute neutrophil counts (ANC), absolute lymphocyte counts (ALC), and neutrophil-to-lymphocyte ratio (NLR), are independent prognostic factors in cancer (1–6). Moreover, NLR could be addressed as a biomarker for the degree of natural and adaptive immune response against tumor (7–11).

While these leukocyte parameters are systemic biomarkers of host immunity, tumor-infiltrating lymphocytes (TILs) are tissue-based biomarkers for adaptive immune response. TILs, which are directly measured in malignant tissues, have been highlighted as a biomarker for predicting treatment response to chemotherapy in patients with breast cancer (9, 12–17). The level of TILs is directly influenced by the host immune system because it is a product of tumor-host immune interactions. Thus, it is worthwhile to explore the relationship between TILs and host immunity. Among the various modalities for assessing host immunity function, peripheral blood cell count is the easiest and most convenient to perform. However, the relationship between TILs and blood cell counts in patients with breast cancer is unclear. Here, we aimed to evaluate the association between TILs and blood cell counts including ALC, ANC, and NLR in treatment-naïve breast cancer.

## II. Material and Methods

### 1. Patients

From August 2016 to July 2018, we enrolled patients with invasive breast cancer who underwent primary breast surgery without preoperative chemotherapy in Gangnam Severance Breast Cancer Center. Those who have (i) a history of transfusion within 3 months, (ii) active infection, (iii) acute/chronic inflammatory disease, and (iv) autoimmune disease were excluded. Clinical data including age at the time of surgery, nuclear grade (NG), histologic grade (HG), tumor size, lymph node status, estrogen receptor (ER) status, progesterone receptor (PR) status, human epidermal growth factor receptor-2 (HER2) status, and Ki-67 labelling index were collected from the medical database. Peripheral blood cell counts were used to determine the ANC, ALC, and NLR. The TNM stage was classified based on the American Joint Committee on Cancer, 7th edition, and tumor grade was determined using the modified Scarf-Bloomer– Richardson grading system.

### 2. Assessment of TIL scores and definition of LPBC

The TIL score was evaluated as previously described (18, 19). A pathologist (YJC) blinded to the clinical information reviewed the histologic features using hematoxylin and eosin (H&E) staining. Briefly, the TIL level was evaluated in treatment-naïve surgical specimens and was scored according to the standardized methodology proposed by the international TIL Working Group (16). Stromal TILs were evaluated according to the guideline. The tumor area, defined by invasive tumor cells, was identified. All mononuclear cells, including lymphocytes and plasma cells but not polymorphonuclear leukocytes, were counted. The area outside the tumor border, around the intraductal component, and normal lobules were excluded. Within the tumor border, areas showing crush artifacts and necrosis were also excluded. For each case, the average TIL was counted on a representative section of the whole tumor, and the average score was reported as a percentage.

In this study, lymphocyte-predominant breast cancer (LPBC) was defined as tumors having high TIL levels (≥50%). The cut-off point of 50% was set according to previous studies using treatment-naïve surgical specimen (5, 13).

### 3. Immunohistochemistry and interpretation

ER (1:100 clone 6F11; Novocastra, Newcastle upon Tyne, UK), PR (clone 16; Novocastra), HER2 (4B5 rabbit monoclonal antibody; Ventana Medical Systems, Tucson, AZ, USA), and Ki-67 (MIB-1;Dako, Glostrup, Denmark) were stained using formalin fixed paraffin-embedded tissue sections as previously described (20, 21). ER positivity and PR expression on immunohistochemistry were defined according to the modified Allred system. In our study, ER and PR positivity were defined as an Allred score ≥ 3. HER2 staining was analyzed according to the American Society of Clinical Oncology/College of American Pathologists guidelines (22). HER2 immunostaining was considered positive when strong membranous staining (3+) was observed, whereas 0 and 1+ were regarded as negative. The cases showing 2+ HER2 expression were further evaluated for HER2 amplification via silver in situ hybridization. Nuclear positivity of Ki-67 staining was evaluated, and the percentage of positive tumor cell (range 0%-100%) was reported as the Ki-67 labeling index. For the breast cancer subtyping, the following definitions were used: i) luminal/HER2(-): ER positive and/or PR positive, and HER2 negative; ii) HER2(+): HER2 positive regardless of ER and PR status; iii) TNBC: ER negative, PR negative, and HER2 negative.

### 4. Complete blood count examination

Blood cell counts including hemoglobin, platelet, WBC, ANC, ALC, neutrophil percentage, lymphocyte percentage, and NLR were obtained from pretreatment laboratory data. Blood samples were collected during the treatment-naïve state after diagnosis. WBC counts were performed at the Department of Laboratory Medicine, Gangnam Severance Hospital using an automated counting machine (Sysmex XN-series; Kobe, Japan). The Wright or Mary-Grinewald-Giemsa technique was used to determine the differential count; briefly, a drop of blood was thinly spread over a glass-slide, air dried, and stained with Romanowsky stain. WBC differentiation was commonly performed using automated fractionation machines, but it was also performed manually in cases of morphologic abnormalities. The neutrophil and lymphocyte counts were calculated as the total WBC multiplied by the percentage of each. Meanwhile, NLR was calculated as ANC divided by ALC.

### 5. Statistical analysis

Pearson’s R was calculated to measure the correlative value between continuous TIL levels and markers of peripheral blood cell counts. Student’s *t*-test or one-way analysis of variation (ANOVA) was used to compare means. Categorical variables were compared using Chi-square test or Fisher’s exact test. Clinico-pathologic factors associated with LPBC were analyzed using the logistic regression analysis. The variables showing a statistical significance in the univariate analysis were entered in the multivariate analysis. All statistical analyses were performed using SPSS version 23 (SPSS; Chicago, IL), and a *p*-value less than 0.05 or odds ratios with a 95% confidence interval (CI) not including 1 was considered significant.

## III. Results

### 1. Baseline characteristics

A total of 684 patients were included in this study. The median patient age was 51 (24-91) years. The mean value and standard deviation of TIL in all patients were 23.95±25.52. The mean values of ANC, ALC, and NLR were 3.6 (K/µL), 1.98 (K/µL), and 1.68, respectively. The patients were divided into the LPBC and non-LPBC groups according to the TIL level. In total, 116 (17.0%) patients had LPBC, and 568 (83.0%) patients had non-LPBC. In Table 1, the LPBC group had higher rates of high NG (*P*<0.001), high HG (*P*<0.001), negative ER (*P*<0.001), negative PR (*P*<0.001), and HER2 overexpression (*P*<0.001). The mean Ki-67 expression was significantly higher in the LPBC group than that in the non-LPBC group (*p*<0.001; Table 1). In addition, the rates of breast cancer subtypes were significantly different between the two groups; the LPBC group had higher rates of HER-positive and TNBC subtypes than the non-LPBC group (*P*<0.001; Fig S1). The LPBC group also had significantly higher Ki-67 labelling index.

**Table 1.** Baseline characteristics according to lymphocyte-predominant breast cancer (LPBC) classification.

|  | LPBC (n=116) | Non-LPBC (n=568) | *P* value |
| --- | --- | --- | --- |
| Age (years) | 52.14±11.02 | 52.57±10.89 | 0.695 |
| ER |  |  | <0.001 |
| Positive | 58 (50) | 498 (87.7) |  |
| Negative | 58 (50) | 70 (12.3) |  |
| PR |  |  | <0.001 |
| Positive | 38 (32.8) | 431 (75.9) |  |
| Negative | 78 (67.2) | 137 (24.1) |  |
| HER2<sup>1,2</sup> |  |  | <0.001 |
| Positive | 55 (47.4) | 82 (14.4) |  |
| Negative | 61 (52.6) | 486 (85.6) |  |
| T stage |  |  | 0.417 |
| I | 76 (65.5) | 396 (69.7) |  |
| II | 38 (32.8) | 154 (27.1) |  |
| III | 2 (1.7) | 18 (3.2) |  |
| N stage |  |  | 0.612 |
| 0 | 97 (83.6) | 446 (78.5) |  |
| I | 17 (14.7) | 98 (17.3) |  |
| II | 1 (0.9) | 16 (2.8) |  |
| III | 1 (0.9) | 8 (1.4) |  |
| TNM Stage* |  |  | 0.405 |
| I | 66 (56.9) | 336 (59.2) |  |
| II | 47 (40.5) | 203 (35.7) |  |
| III | 3 (2.6) | 29 (5.1) |  |
| NG<sup>1</sup> |  |  | <0.001 |
| I and II | 35 (31.3) | 449 (79.8) |  |
| III | 77 (68.8) | 114 (20.2) |  |
| HG<sup>1</sup> |  |  | <0.001 |
| I and II | 57 (55.3) | 496 (89.4) |  |
| III | 46 (44.7) | 59 (10.6) |  |
| Ki-67<sup>1</sup> (%) | 38.90±25.77 | 14.61±18.05 | <0.001 |

| Subtypes |  |  | <0.001 |
| --- | --- | --- | --- |
| Luminal/HER2(-) | 32 (27.6) | 447 (78.7) |  |
| HER2(+) | 55 (47.4) | 82 (14.4) |  |
| TNBC | 29 (25) | 39 (6.9) |  |

Data are presented as n (%) or mean±SD.
Abbreviations: ER, estrogen receptor; PR, progesterone receptor; HER-2, human epidermal growth factor receptor-2; NG, nuclear grade; HG, histologic grade; TNBC, triple negative breast cancer
\*Staging was according to the AJCC 7<sup>th</sup> edition.
<sup>1</sup>Missing value
<sup>2</sup>HER-2 positive was defined by 3 positive on immunohistochemistry or amplification on fluorescence *in situ* hybridization

In addition, the mean ANC of the LPBC group was significantly higher than that of the non-LPBC (*P*=0.003; Fig 1A). However, means of ALC and NLR did not differ between two groups (Fig 1B, *P*=0.677; Fig 1C, *P*=0.196, respectively).

**Fig 1.**
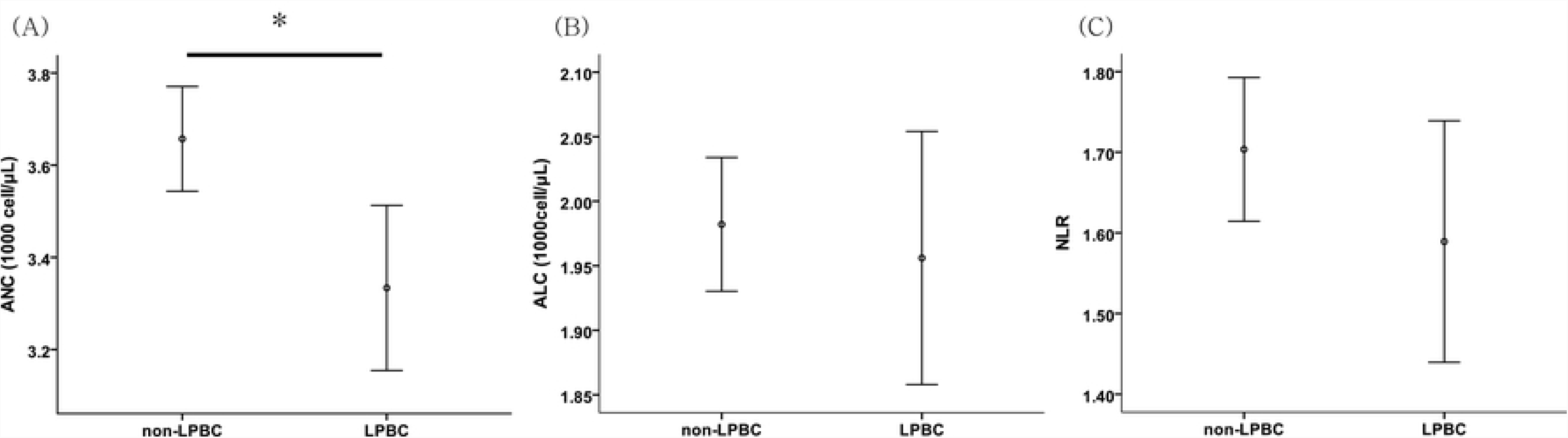
Comparison of peripheral blood markers between the LPBC and the non-LPBC group. (A) The mean ANC of the LPBC group was significantly lower than that of the non-LPBC group (3.33 vs 3.66 K/μL, * *P*=0.003, Student’s *t*-test). However, the mean ALC (B) and NLR (C) did not differ between the two groups (ALC: 1.96 vs 1.98 K/μL, *P*=0.677; NLR: 1.59 vs 1.70, *P*=0.196; Student’s *t*-test).

### 2. Association between TILs and blood cell markers

We compared the means of the blood cell markers among the subtypes because LPBC was more frequent in HER2-positive or TNBC subtypes (Fig S1). However, there were no significant differences in the means of three blood cell markers (ANC, ALC, and NLR) between the cancer subtypes (Fig S2).

Pearson’s R test was performed to identify the correlation between TIL levels and blood cell markers. A moderate inverse correlation without a statistical significance was observed between TIL levels and ANC (Pearson’s R= −0.50, *P*=0.192; Fig S3). We also found no significant correlation between continuous TIL levels and two blood markers (ALC and NLR; Fig S4).

### 3. Logistic regression analysis for LPBC

In the univariate analysis for the clinico-pathologic factors associated with LPBC, the significant variables were ANC, breast cancer subtype, Ki-67, NG, and HG. Meanwhile, continuous ANC remained an independent variable associated with LPBC in multivariate analysis (OR: 0.736, 95% CI: 0.591-0.917, *P*=0.006; Table 2). The subtypes, Ki-67 expression, and NG were also found to be significant variables in identifying LPBC.

**Table 2.** Odds ratios (ORs) and 95% confidential intervals (CIs) of the blood cell counts for identifying LPBC.

|  | Univariate analysis |  | Multivariate analysis |  |
| --- | --- | --- | --- | --- |
|  | ORs (95% CIs) | *P* value | ORs (95% CIs) | *P* value |
| ANC | 0.809 (0.681-0.962) | 0.016 | 0.736 (0.591-0.917) | 0.006 |
| ALC | 0.932 (0.669-1.298) | 0.676 |  |  |
| NLR | 0.792 (0.618-1.015) | 0.065 |  |  |
| Subtype |  | <0.001 |  | 0.001 |
| Luminal/HER2(-) | Ref |  | Ref |  |
| HER2(+) | 9.639 (5.709-15.375) |  | 3.215 (1.755-5.890) |  |
| TNBC | 10.387 (5.702-18.920) |  | 2.494 (1.092-5.698) |  |
| Ki-67 | 1.043 (1.034-1.052) | <0.001 | 1.028 (1.015-1.042) | <0.001 |
| NG |  | <0.001 |  | <0.001 |
| I and II | Ref |  | Ref |  |
| III | 2.944 (2.351-3.685) |  | 3.072 (1.662-5.677) |  |
| HG |  | <0.001 |  | 0.646 |
| I and II | Ref |  | Ref |  |
| III | 2.605 (2.056-3.300) |  | 0.844 (0.409-1.742) |  |
| T stage |  | 0.368 |  |  |
| I | Ref |  |  |  |
| II | 1.286 (0.835-1.980) |  |  |  |
| III | 0.579 (0.132-2.547) |  |  |  |
| N stage |  | 0.528 |  |  |
| 0 | Ref |  |  |  |
| I | 0.798 (0.456-1.396) |  |  |  |
| II | 0.287 (0.038-2.193) |  |  |  |
| III | 0.575 (0.071-4.649) |  |  |  |
| AJCC stage |  | 0.380 |  |  |
| I | Ref |  |  |  |
| II | 1.179 (0.780-1.781) |  |  |  |
| III | 0.527 (0.156-1.780) |  |  |  |

## IV. Discussion

In this study, we investigated whether peripheral blood cell counts are associated with TILs in breast cancer. We hypothesized that TILs are associated with blood cell counts, which reflect local and host immunity, respectively. To the best of our knowledge, this is the first study investigating the relationship between TILs and blood cell markers. We found a significantly reduced mean ANC in those with LPBC compared to those with non-LPBC. Moreover, continuous ANC was a significant predictive factor of LPBC, independent of the cancer subtypes.

The negative correlation observed between high ANC and LPBC in our study is supported by the fact that neutrophils may act against the immune system via several mechanisms. Experimental data suggested that neutrophils could suppress the cytolytic activity of lymphocytes, natural killer cells, and activated T-cells when it is co-cultured with lymphocytes form normal healthy donor. Moreover, activated neutrophils have been reported to secrete myeloperoxidase, resulting in the suppression of lymphocyte function (7).

Moreover, tumor-associated neutrophils might influence local tumor immunity and tumor progression by regulating the tumor microenvironment. The enzymatic activity of neutrophils has been found to promote remodeling of the extracellular matrix, which results in the release of basic fibroblast growth factor and migration of either endothelial cells or tumor cells. Also, reactive oxygen species released by neutrophils further decrease the adhesive propensity of extracellular matrix and inhibit apoptosis of the tumor cells via activation of nuclear factor-κB (7, 8). The modulated tumor microenvironment might contribute to tumor growth and acquisition of metastatic capability. Specifically, neutrophil-derived oncostatin M stimulates cancer cells to secrete vascular endothelial growth factor and increases invasiveness in breast cancer (23).

We also found that LPBC was associated with the cancer subtypes; specifically, the rate of LPBC was higher in the HER2 and TNBC subtypes are than that in the luminal/HER2-negative subtype. Another notable finding was that the Ki-67 labelling index was correlated with LPBC, which may be explained by the fact that tumors with high Ki-67 labelling index were more frequent in the HER2 or TNBC subtypes. These findings provide evidence that our data is reliable.

Clinically, a previous study showed that high ANC could be a poor prognostic marker in patients with breast cancer (24), supporting our findings that high ANC may negatively impact TILs in breast cancer. The association between NLR and TILs has not been identified, although NLR is a well-known poor prognostic marker in various cancers including breast cancer (9, 25–30). The relationship between NLR and TILs warrants further research.

Interestingly, emerging evidence suggests that high ANC could be a negative predictor of response to immune checkpoint inhibitors (ICI) for cancer such as lung cancer and melanoma (31–33). By contrast, high TILs are positive predictors of treatment response to ICI (34, 35). Contrasting responses to ICI according to high ANC and TILs indirectly support our results, but further studies are still needed to clearly establish the predictive value of TILs in the treatment response to ICI.

This study also has some limitations, including the absence of survival analyses due to short-term follow-up. Assessment of the clinical outcomes of the study population might have helped to specify the prognostic capability of these biomarkers. Another limitation is that neutrophil and lymphocyte counts may vary according to individual physiological and pathological processes such as infectious condition. The association between NLR and TILs in breast cancer warrants further research. Despite these limitations, we found relevant evidence showing specific correlations between TILs and ANC, providing scientific insight on the host-tumor immune interaction in breast cancer.

## V. Conclusion

Low peripheral ANC might be correlated with LPBC, supporting the hypothesis that systemic immune cell counts might be associated with the tumor-immune microenvironment. A possible association between peripheral ANC and tumor-associated neutrophil reflecting the tumor microenvironment should be studied.

## Acknowledgements

We acknowledge all authors in our paper.

## Supporting information

**S1 Fig. Proportion of breast cancer subtypes according to LPBC classification. The** LPBC group had higher rates of HER2-positive and TNBC subtypes than the non-LPBC group (*P*<0.001, ANOVA)

**S2 Fig. Comparison of peripheral blood markers according to breast cancer subtypes.** There were no significant differences in peripheral blood markers according to breast cancer subtypes. (A) ANC: 3634 vs 3413 vs 3756 cells/μL, *P=*0.136. (B) ALC: 1976 vs 2003 vs 1940 cells/μL, *P*=0.781. (C) NLR: 1.70 vs 1.57 vs 1.83, *P*=0.202; ANOVA.

**S3 Fig. Pearson correlation analysis for the association between TIL and ANC.** The correlation coefficient value (Pearson’s R) was −0.50, indicating moderate inverse correlation without statistical significance (*P*=0.192).

**S4 Fig. Pearson correlation analysis for the association between TIL and other peripheral blood markers. (A) ALC and (B) NLR**. The correlation coefficient value (Pearson’s R) of ALC and NLR were −0.09 and 0.009, respectively, and there were no significant correlations (ALC: *P*=0.815; NLR: *P*=0.816).

## Reference

1. Adams S, Gray RJ, Demaria S, Goldstein L, Perez EA, Shulman LN, et al. Prognostic value of tumor-infiltrating lymphocytes in triple-negative breast cancers from two phase III randomized adjuvant breast cancer trials: ECOG 2197 and ECOG 1199. J Clin Oncol. 2014;32(27):2959–66.

2. Denkert C, Loibl S, Noske A, Roller M, Muller BM, Komor M, et al. Tumor-associated lymphocytes as an independent predictor of response to neoadjuvant chemotherapy in breast cancer. J Clin Oncol. 2010;28(1):105–13.

3. Denkert C, von Minckwitz G, Brase JC, Sinn BV, Gade S, Kronenwett R, et al. Tumor-infiltrating lymphocytes and response to neoadjuvant chemotherapy with or without carboplatin in human epidermal growth factor receptor 2-positive and triple-negative primary breast cancers. J Clin Oncol. 2015;33(9):983–91.

4. Lee HJ, Seo JY, Ahn JH, Ahn SH, Gong G. Tumor-associated lymphocytes predict response to neoadjuvant chemotherapy in breast cancer patients. J Breast Cancer. 2013;16(1):32–9.

5. Loi S, Sirtaine N, Piette F, Salgado R, Viale G, Van Eenoo F, et al. Prognostic and predictive value of tumor-infiltrating lymphocytes in a phase III randomized adjuvant breast cancer trial in node-positive breast cancer comparing the addition of docetaxel to doxorubicin with doxorubicin-based chemotherapy: BIG 02-98. J Clin Oncol. 2013;31(7):860–7.

6. Seo AN, Lee HJ, Kim EJ, Kim HJ, Jang MH, Lee HE, et al. Tumour-infiltrating CD8+ lymphocytes as an independent predictive factor for pathological complete response to primary systemic therapy in breast cancer. Br J Cancer. 2013;109(10):2705–13.

7. De Larco JE, Wuertz BR, Furcht LT. The potential role of neutrophils in promoting the metastatic phenotype of tumors releasing interleukin-8. Clinical cancer research: an official journal of the American Association for Cancer Research. 2004;10(15):4895–900.

8. el-Hag A, Clark RA. Immunosuppression by activated human neutrophils. Dependence on the myeloperoxidase system. Journal of immunology (Baltimore, Md: 1950). 1987;139(7):2406–13.

9. Ethier JL, Desautels D, Templeton A, Shah PS, Amir E. Prognostic role of neutrophil-to-lymphocyte ratio in breast cancer: a systematic review and meta-analysis. Breast cancer research: BCR. 2017;19(1):2.

10. Sacdalan DB, Lucero JA, Sacdalan DL. Prognostic utility of baseline neutrophil-to-lymphocyte ratio in patients receiving immune checkpoint inhibitors: a review and meta-analysis. OncoTargets and therapy. 2018;11:955–65.

11. Labomascus S, Fughhi I, Bonomi P, Fidler MJ, Borgia JA, Basu S, et al. Neutrophil to lymphocyte ratio as predictive of prolonged progression free survival (PFS) and overall survival (OS) in patients with metastatic non-small cell lung cancer (NSCLC) treated with second-line PD-1 immune checkpoint inhibitors. Journal of Clinical Oncology. 2017;35(15_suppl):e14530–e.

12. Can C, Baseskioglu B, Yilmaz M, Colak E, Ozen A, Yenilmez A. Pretreatment parameters obtained from peripheral blood sample predicts invasiveness of bladder carcinoma. Urologia internationalis. 2012;89(4):468–72.

13. Dieci MV, Mathieu MC, Guarneri V, Conte P, Delaloge S, Andre F, et al. Prognostic and predictive value of tumor-infiltrating lymphocytes in two phase III randomized adjuvant breast cancer trials. Annals of oncology: official journal of the European Society for Medical Oncology. 2015;26(8):1698–704.

14. Gutkin DW, Shurin MR. Clinical evaluation of systemic and local immune responses in cancer: time for integration. Cancer immunology, immunotherapy: CII. 2014;63(1):45–57.

15. Moore MM, Chua W, Charles KA, Clarke SJ. Inflammation and cancer: causes and consequences. Clinical pharmacology and therapeutics. 2010;87(4):504–8.

16. Salgado R, Denkert C, Demaria S, Sirtaine N, Klauschen F, Pruneri G, et al. The evaluation of tumor-infiltrating lymphocytes (TILs) in breast cancer: recommendations by an International TILs Working Group 2014. Annals of oncology: official journal of the European Society for Medical Oncology. 2015;26(2):259–71.

17. West NR, Kost SE, Martin SD, Milne K, Deleeuw RJ, Nelson BH, et al. Tumour-infiltrating FOXP3(+) lymphocytes are associated with cytotoxic immune responses and good clinical outcome in oestrogen receptor-negative breast cancer. Br J Cancer. 2013;108(1):155–62.

18. Ahn SG, Cha YJ, Bae SJ, Yoon C, Lee HW, Jeong J. Comparisons of tumor-infiltrating lymphocyte levels and the 21-gene recurrence score in ER-positive/HER2-negative breast cancer. BMC cancer. 2018;18(1):320.

19. Cha YJ, Ahn SG, Bae SJ, Yoon CI, Seo J, Jung WH, et al. Comparison of tumor-infiltrating lymphocytes of breast cancer in core needle biopsies and resected specimens: a retrospective analysis. Breast cancer research and treatment. 2018;171(2):295–302.

20. Ahn SG, Dong SM, Oshima A, Kim WH, Lee HM, Lee SA, et al. LOXL2 expression is associated with invasiveness and negatively influences survival in breast cancer patients. Breast cancer research and treatment. 2013;141(1):89–99.

21. Harvey JM, Clark GM, Osborne CK, Allred DC. Estrogen receptor status by immunohistochemistry is superior to the ligand-binding assay for predicting response to adjuvant endocrine therapy in breast cancer. J Clin Oncol. 1999;17(5):1474–81.

22. Wolff AC, Hammond ME, Schwartz JN, Hagerty KL, Allred DC, Cote RJ, et al. American Society of Clinical Oncology/College of American Pathologists guideline recommendations for human epidermal growth factor receptor 2 testing in breast cancer. J Clin Oncol. 2007;25(1):118–45.

23. Queen MM, Ryan RE, Holzer RG, Keller-Peck CR, Jorcyk CL. Breast cancer cells stimulate neutrophils to produce oncostatin M: potential implications for tumor progression. Cancer research. 2005;65(19):8896–904.

24. Wariss BR, de Souza Abrahao K, de Aguiar SS, Bergmann A, Thuler LCS. Effectiveness of four inflammatory markers in predicting prognosis in 2374 women with breast cancer. Maturitas. 2017;101:51–6.

25. Beal EW, Wei L, Ethun CG, Black SM, Dillhoff M, Salem A, et al. Elevated NLR in gallbladder cancer and cholangiocarcinoma - making bad cancers even worse: results from the US Extrahepatic Biliary Malignancy Consortium. HPB: the official journal of the International Hepato Pancreato Biliary Association. 2016;18(11):950–7.

26. Huang QT, Zhou L, Zeng WJ, Ma QQ, Wang W, Zhong M, et al. Prognostic Significance of Neutrophil-to-Lymphocyte Ratio in Ovarian Cancer: A Systematic Review and Meta-Analysis of Observational Studies. Cellular physiology and biochemistry: international journal of experimental cellular physiology, biochemistry, and pharmacology. 2017;41(6):2411–8.

27. Marchioni M, Primiceri G, Ingrosso M, Filograna R, Castellan P, De Francesco P, et al. The Clinical Use of the Neutrophil to Lymphocyte Ratio (NLR) in Urothelial Cancer: A Systematic Review. Clinical genitourinary cancer. 2016;14(6):473–84.

28. Mei Z, Shi L, Wang B, Yang J, Xiao Z, Du P, et al. Prognostic role of pretreatment blood neutrophil-to-lymphocyte ratio in advanced cancer survivors: A systematic review and meta-analysis of 66 cohort studies. Cancer treatment reviews. 2017;58:1–13.

29. Wei B, Yao M, Xing C, Wang W, Yao J, Hong Y, et al. The neutrophil lymphocyte ratio is associated with breast cancer prognosis: an updated systematic review and meta-analysis. OncoTargets and therapy. 2016;9:5567–75.

30. Yang JJ, Hu ZG, Shi WX, Deng T, He SQ, Yuan SG. Prognostic significance of neutrophil to lymphocyte ratio in pancreatic cancer: a meta-analysis. World journal of gastroenterology. 2015;21(9):2807–15.

31. Okuhira H, Yamamoto Y, Inaba Y, Kunimoto K, Mikita N, Ikeda T, et al. Prognostic factors of daily blood examination for advanced melanoma patients treated with nivolumab. Bioscience trends. 2018;12(4):412–8.

32. Soda H, Ogawara D, Fukuda Y, Tomono H, Okuno D, Koga S, et al. Dynamics of blood neutrophil-related indices during nivolumab treatment may be associated with response to salvage chemotherapy for non-small cell lung cancer: A hypothesis-generating study. Thoracic cancer. 2018.

33. Zer A, Sung MR, Walia P, Khoja L, Maganti M, Labbe C, et al. Correlation of Neutrophil to Lymphocyte Ratio and Absolute Neutrophil Count With Outcomes With PD-1 Axis Inhibitors in Patients With Advanced Non-Small-Cell Lung Cancer. Clinical lung cancer. 2018;19(5):426–34.e1.

34. Loi S, Adams S, Schmid P, Cortés J, Cescon DW, Winer EP, et al. LBA13Relationship between tumor infiltrating lymphocyte (TIL) levels and response to pembrolizumab (pembro) in metastatic triple-negative breast cancer (mTNBC): Results from KEYNOTE-086. Annals of Oncology. 2017;28(suppl_5).

35. Tumeh PC, Hellmann MD, Hamid O, Tsai KK, Loo KL, Gubens MA, et al. Liver Metastasis and Treatment Outcome with Anti-PD-1 Monoclonal Antibody in Patients with Melanoma and NSCLC. Cancer immunology research. 2017;5(5):417–24.

